# Polymorphic inversions underlie the shared genetic susceptibility to prevalent common diseases

**DOI:** 10.1101/859280

**Authors:** Juan R González, Carlos Ruiz-Arenas, Alejandro Cáceres, Ignasi Morán, Marcos López, Lorena Alonso, Ignacio Tolosana, Marta Guindo-Martínez, Josep M Mercader, Tonu Esko, David Torrents, Josefa González, Luis A Pérez-Jurado

**Author notes:** **Corresponding author:** Juan R González, Bioinformatics Research Group in Epidemiology, Barcelona Institute for Global Health (ISGlobal), Avd. Dr Aiguader, 88, 08003 Barcelona, Spain.

## Abstract

The burden of several common diseases including obesity, diabetes, hypertension, asthma, and depression is increasing in most world populations. However, the mechanisms underlying the numerous epidemiological and genetic correlations among these disorders remain largely unknown. We investigated whether common polymorphic inversions underlie the shared genetic influence of these disorders. We performed the largest inversion association analysis to date, including 21 inversions and 25 obesity-related traits, on a total of 408,898 Europeans, and validated the results in 67,299 independent individuals. Seven inversions were associated with multiple diseases while inversions at 8p23.1, 16p11.2 and 11q13.2 were strongly associated with the co-occurrence of obesity with other common diseases. Transcriptome analysis across numerous tissues revealed strong candidate genes of obesity-related traits. Analyses in human pancreatic islets indicated the potential mechanism of inversions in the susceptibility of diabetes by disrupting the cis-regulatory effect of SNPs from their target genes. Our data underscore the role of inversions as major genetic contributors to the joint susceptibility to common complex diseases.

---

Obesity is a disorder with increasing but non-uniform prevalence in the world population and one of the major public health burdens^1^. Obesity derived morbidity and years of life lost strongly associate to a broad range of highly prevalent diseases, including type 2 diabetes, cardiovascular disease, asthma and psychological disturbance such as depression, among others^2^. While the causes underlying the multiple co-occurrences of obesity are likely complex and diverse, common mechanisms underlying these comorbidities, which are potential targets for preventive or therapeutic intervention, are largely unknown.

Genomic inversions, copy-neutral changes in the orientation of chromosomal segments with respect to the reference, are excellent candidates for being important contributors to the genetic architecture of common diseases. Inversion polymorphisms can alter the function of the including and neighboring genes by multiple mechanisms, disrupting genes, separating their regulatory elements, affecting chromatin structure, and maintaining a strong linkage of functional variants within an interval that escape recombination. Therefore, by putatively affecting multiple genes in numerous ways, inversions are important sources of shared genomic variation underlying different human diseases and traits. Consequently, human inversions show genetic influences in multiple phenotypes. For instance, the common inversion at 8p23.1 has been independently linked to obesity^3^, autism^4^, neuroticism^5^ and several risk behavior traits^6^, while inversion at 17q21.31 has been associated with Alzheimer^7^ and Parkinson^8^ diseases, heart failure^9^ and intracranial volume^10^. We previously reported a ~40% of population attributable risk for the co-occurrence of asthma and obesity given by a common inversion polymorphism at 16p11.2^11^. In addition, transcriptional effects have been documented in several tissues for inversions at 17q21.31^8,12^, and 16p11.2^11^.

It has been estimated that each human genome contains about 156 inversions^13^. Therefore, inversions constitute a substantial source of genetic variability. Many of those polymorphic inversions show signatures of positive or balancing selection associated with functional effects^14^. However, the overall impact of polymorphic inversions on human health remains largely unknown because they are difficult to genotype in large cohorts. We overcame this limitation by recently reporting a subset of 20 inversions that can be genotyped with SNP array data as they are old in origin, low or not recurrent and frequent in the population^15^. We have also included an additional inversion in our catalog, 16p11.2, previously validated and genotyped in diverse populations^11^. Three of the inversions are submicroscopic (0.45-4 Mb), flanked by large segmental duplications and contain multiple genes. Five are small (0.7-5 Kb) and intragenic, and 13 are intergenic of variable size (0.7-90 Kb) but highly enriched in pleiotropic genomic regions^16^. While this is clearly not a comprehensive set of inversions, it is probably the largest set that can be genotyped in publicly available datasets.

In this manuscript, we aimed to study the association of 21 common polymorphic inversions in Europeans with highly prevalent co-morbid disorders and related traits. We particularly aimed to decipher the role of inversions in known epidemiological co-occurrences with obesity such as diabetes, hypertension, asthma and mental diseases like depression, bipolar disorder or neuroticism. For significant associations, we investigated whether causal pathways could be established and the most likely underlying mechanisms.

## RESULTS

### Frequency and stratification of inversions in European populations

Using *scoreInvHap,* we first called the inversion status of individuals from the UK Biobank (UKB) with European ancestry (n=408,898). We confirmed the previously reported frequency in the 1000 Genome project of the 21 inversions analyzed in this work (**Table 1**). As inversion frequencies have a strong demographic effect, we also analyzed 12 European countries from the POPRES study (**Supplementary Figure S1).** We observed significant clines along north-south latitude for several inversions (**Table 1 and Supplementary Figure S2A**) as well as subtle ancestral differences (**Supplementary Figure S2B**). Thus, population stratification was considered when performing association analyses as explained in methods section.

**Table 1.**
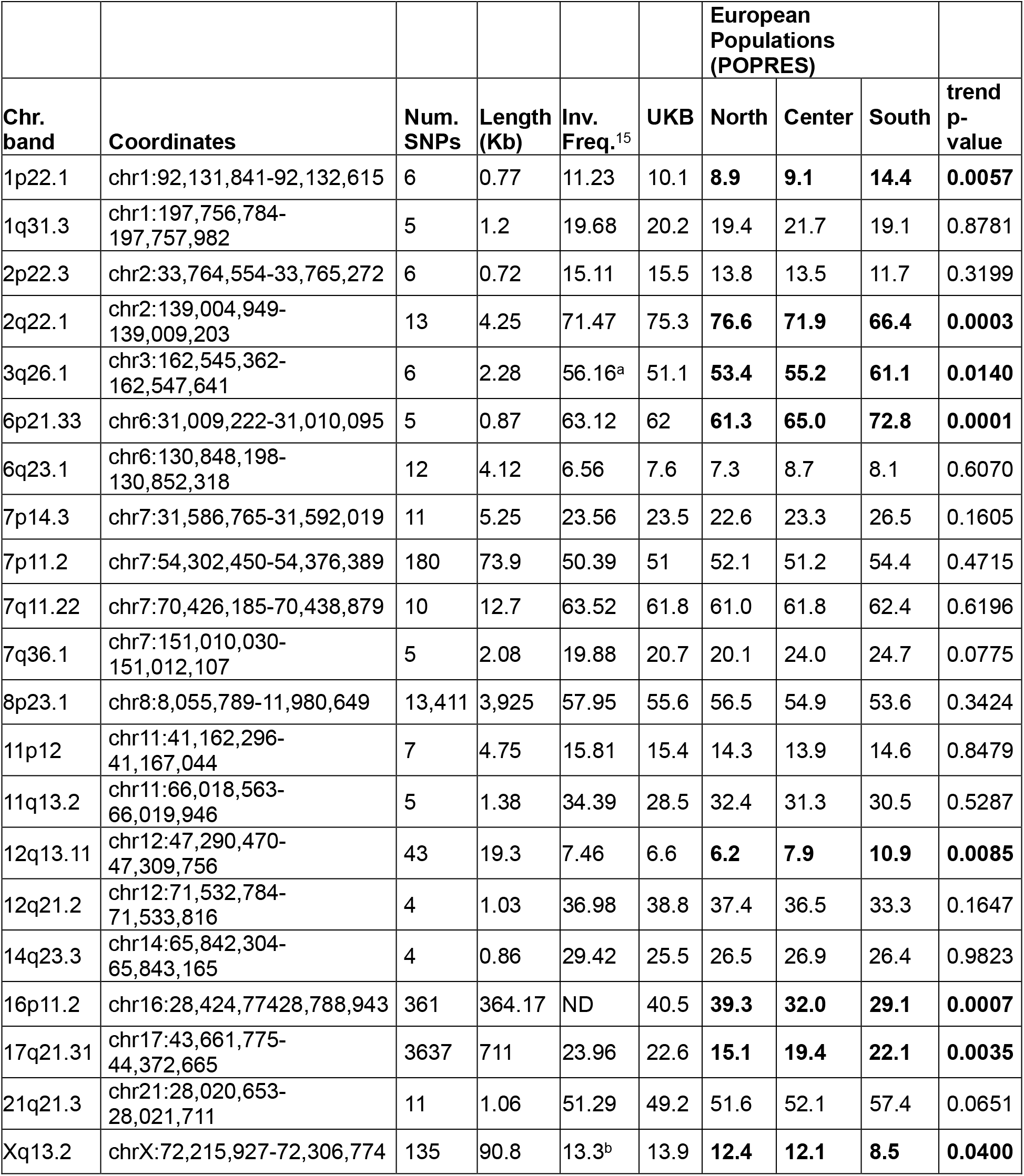
Characteristics of the 21 genomic inversions. The table shows the coordinates, SNP content, size, and inversion frequency obtained from 1000 Genomes as described in Ruiz-Arenas et al^15^, the UKB and European regions (north, center and south) using the regions described in the POPRES dataset (see **Methods**). The p-value correspond to a trend test to assess north-south linear association (in bold those significant at 5% level).

### Inversions at 8p23.1, 16p11.2 robustly associate with obesity and obesity-related traits

The discovery phase of the study used data from UKB. We performed association analyses between the 21 inversions with obesity and co-morbid diseases and traits (see **Methods**). These include obesity, diabetes, stroke, hypertension, asthma, chronic obstructive pulmonary disease (COPD), depression and bipolar disorder, along with related traits or phenotypes classified as morphometric (4 traits), metabolic (5 traits), lipidic (2 traits), respiratory (3 traits) and behavioral (3 traits) (**Figure 1**). **Supplementary Table S1** shows the total number of cases and controls used to perform the association analyses on each trait. The significant associations were further validated in GERA and 70KT2D independent datasets (see **Methods**).

**Figure 1.**
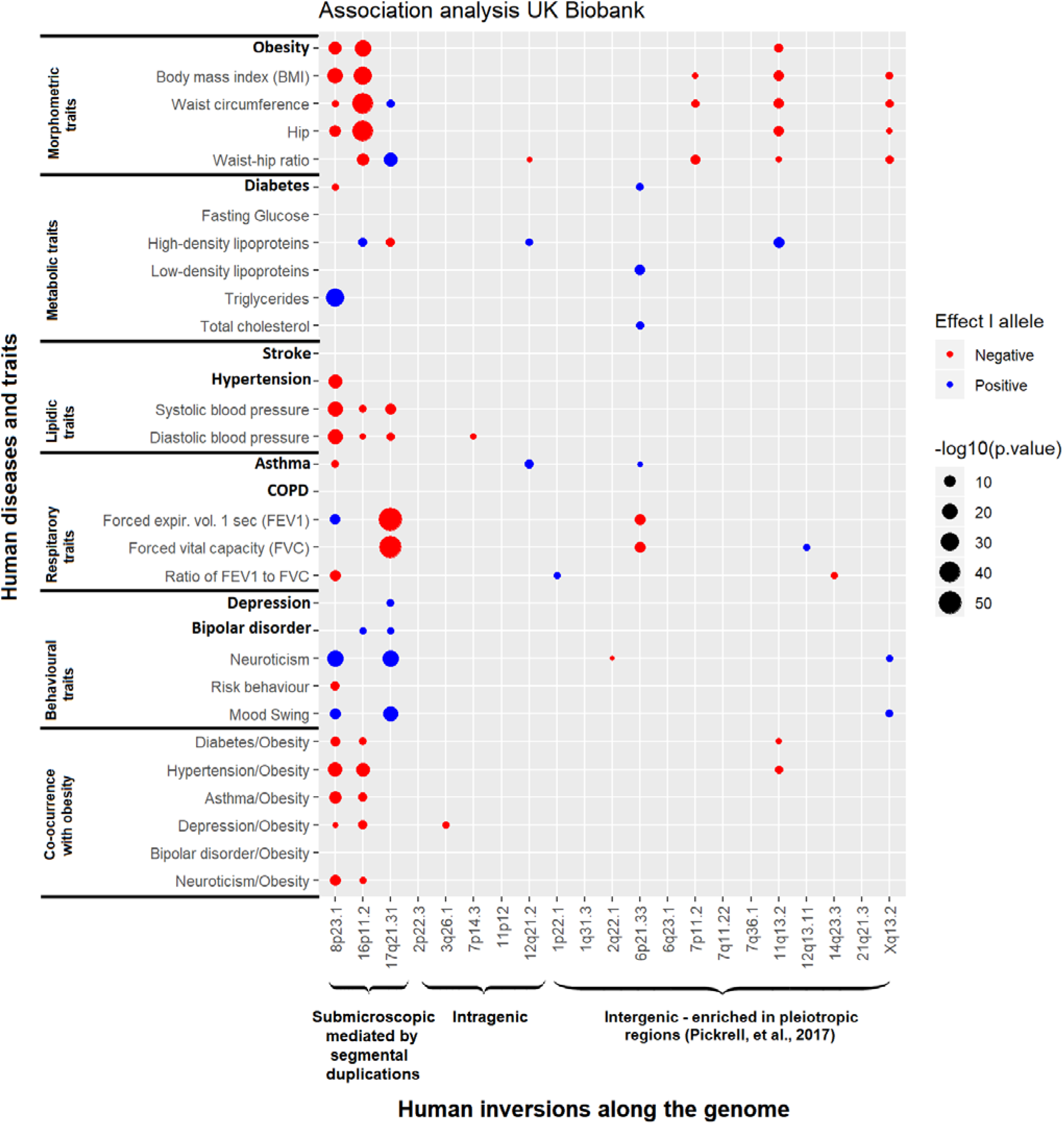
Association analyses between 21 inversions and 8 diseases (in bold) and 17 traits and the co-occurrence of obesity with 6 other complex diseases. Circles represent the direction (color) and the strength (size) of the association for different groups of traits (morphometric, metabolic, lipidic, respiratory and behavioral) and the epidemiological well-established co-occurrence of obesity-related diseases. Inversions are grouped by size and features: 1) submicroscopic are large (0.4-4Mb) encompassing multiple genes and flanked by segmental duplications; 2) intragenic are located within a gene, either intronic or containing one exon; and 3) intergenic are enriched in pleitropic regions

The analyses on the UKB revealed several genetic influences of inversions on obesity and related common diseases (**Figure 1**). We observed a total of 78 significant associations after correcting for the number of inversions analyzed. In general, we observed higher numbers of associations and stronger effects for the largest inversions at 8p23.1, 16p11.2 and 17q21.31, consistent with the fact that they encapsulate more genes. Some smaller inversions such as the ones at 11q13.2 and Xq13.2 also showed notable effects such as shared susceptibility and strength, respectively. We found a prominent inflation of association suggesting common genetic influences of the inversions across multiple phenotypes (**Supplementary Figure S3A**). Some of the associations found have already been reported such as those at inversion 8p23.1 with obesity^3^ and neuroticism^5^ and the one with inversion 16p11.2 with obesity^11^.

As a summary of the novel findings, we observe that inversions at 8p23.1, 16p11.2 and 11q13.2 are all strongly associated with several obesity-related diseases (**Figure 1**). Remarkably, the non-inverted (N) allele of inversion 8p23.1 (i.e. the risk allele) is independently associated with diabetes (OR=1.04, p=1.1×10^−3^), hypertension (OR=1.04, p=7.0×10^−16^) and asthma (OR=1.03, p=7.0×10^−5^) (**Table 2**). The association with diabetes was replicated in the 70KT2D study (**Figure 2A)** (OR =1.08, p=1.1×10^−8^) as well as the association with obesity (OR=1.08, p=5.6×10^−6^) and the association with hypertension which was validated in the GERA study (OR=1.03, p=0.0183) (**Table 2**). We also found a novel significant association between the non-inverted (N) allele of inversion 16p11.2 and obesity (OR=1.05, p=3.9×10^−24^) that was replicated in GERA study (OR=1.07, p=1.4×10^−4^). The significant associations found in the UKB for the inversion 11q13.2 was marginally validated in the GERA study (OR=1.03, p= 0.0712). Consistently, the analysis of UKB study also revealed association of inversions at 8p23.1, 16p11.2 with different obesity-related traits such as body mass index (BMI), waist circumference, high density lipoprotein (HDL) or systolic and diastolic blood pressure among others (**Figure 1**).

**Table 2:** Association between inversions 8p23.1 and 16p1.2 and different obesity-related traits in UKB and replication data sets. The table shows the odds ratios (OR) and their confidence intervals at 95% (CI95%) for the inverted allele and different diseases and the joint co-occurrence with obesity at UKB and replication datasets. The p corresponds to the best genetic model depict in the first column of each inversion.

|  | **Inversion 8p23.1 (effect of risk-Haplotype: N-allele)** |  |  |  |  | **Inversion 16p11.2 (effect of risk-Haplotype: N-allele)** |  |  |  |
| --- | --- | --- | --- | --- | --- | --- | --- | --- | --- |
|  | **UKB** |  | **Replication** |  |  | **UKB** |  | **Replication** |  |
| **Disease** | **OR CI95%** | **p-value** | **OR CI95%** | **p-value** |  | **OR CI95%** | **p-value** | **OR CI95%** | **p-value** |
| **Obesity** | 1.04 (1.03-1.05) | 2.4x10<sup>-13</sup> | 1.08 (1.04-1.11) | 5.6x10<sup>-6</sup> |  | 1.05 (1.04-1.06) | 3.9x10<sup>-24</sup> | 1.07 (1.03-1.10) | 1.4x10<sup>-4</sup> |
| **Diabetes** | 1.04 (1.01-1.06) | 1.1x10<sup>-3</sup> | 1.08 (1.05-1.11) | 1.1x10<sup>-8</sup> |  | 1.02 (0.99-1.04) | 0.1450 | 1.07 (1.04- 1.11) | 1.2x10<sup>-6</sup> |
| **Hypertension** | 1.04 (1.03-1.05) | 7.0x10<sup>-16</sup> | 1.03 (1.00-1.05) | 0.0183 |  | 1.01 (1.00-1.02) | 0.0184 | 1.02 (0.99-1.05) | 0.2127 |
| **Asthma** | 1.03 (1.01-1.04) | 7.0x10<sup>-5</sup> | 1.02 (0.90-1.05) | 0.2225 |  | 1.00 (0.99-1.01) | 0.9529 | 1.00 (0.97-1.04) | 0.8074 |
| **Depression** | 0.98 (0.97-0.99) | 0.0119 | 1.01 (0.97-1.05) | 0.6630 |  | 0.98 (0.96-1.00) | 0.0184 | 1.01 (0.98-1.05) | 0.5384 |
| **Joint occurrence of Obesity with:** |  |  |  |  |  |  |  |  |  |
| Diabetes | 1.08 (1.05-1.11) | 3.1x10<sup>-7</sup> | 1.17 (1.12-1.22) | 1.4 x10<sup>-13</sup> |  | 1.06 (1.03-1.08) | 7.5x10<sup>-5</sup> | 1.13 (1.08- 1.17) | 1.2x10<sup>-8</sup> |
| Hypertension | 1.07 (1.05-1.08) | 1.7x10<sup>-16</sup> | 1.06 (1.02-1.11) | 6.9x10<sup>-3</sup> |  | 1.06 (1.05-1.07) | 2.7x10<sup>-14</sup> | 1.05 (1.00-1.10) | 0.0357 |
| Asthma | 1.08 (1.06-1.10) | 3.0x10<sup>-11</sup> | 1.09 (1.02-1.16) | 9.7x10<sup>-3</sup> |  | 1.05 (1.03-1.07) | 7.4x10<sup>-6</sup> | 1.08 (1.01-1.15) | 0.0287 |
| Depression | 1.04 (1.02-1.07) | 1.4x10<sup>-3</sup> | 1.12 (1.04-1.20) | 3.8x10<sup>-3</sup> |  | 1.06 (1.03-1.08) | 1.4x10<sup>-6</sup> | 1.03 (0.95-1.11) | 0.5241 |

**Figure 2.**
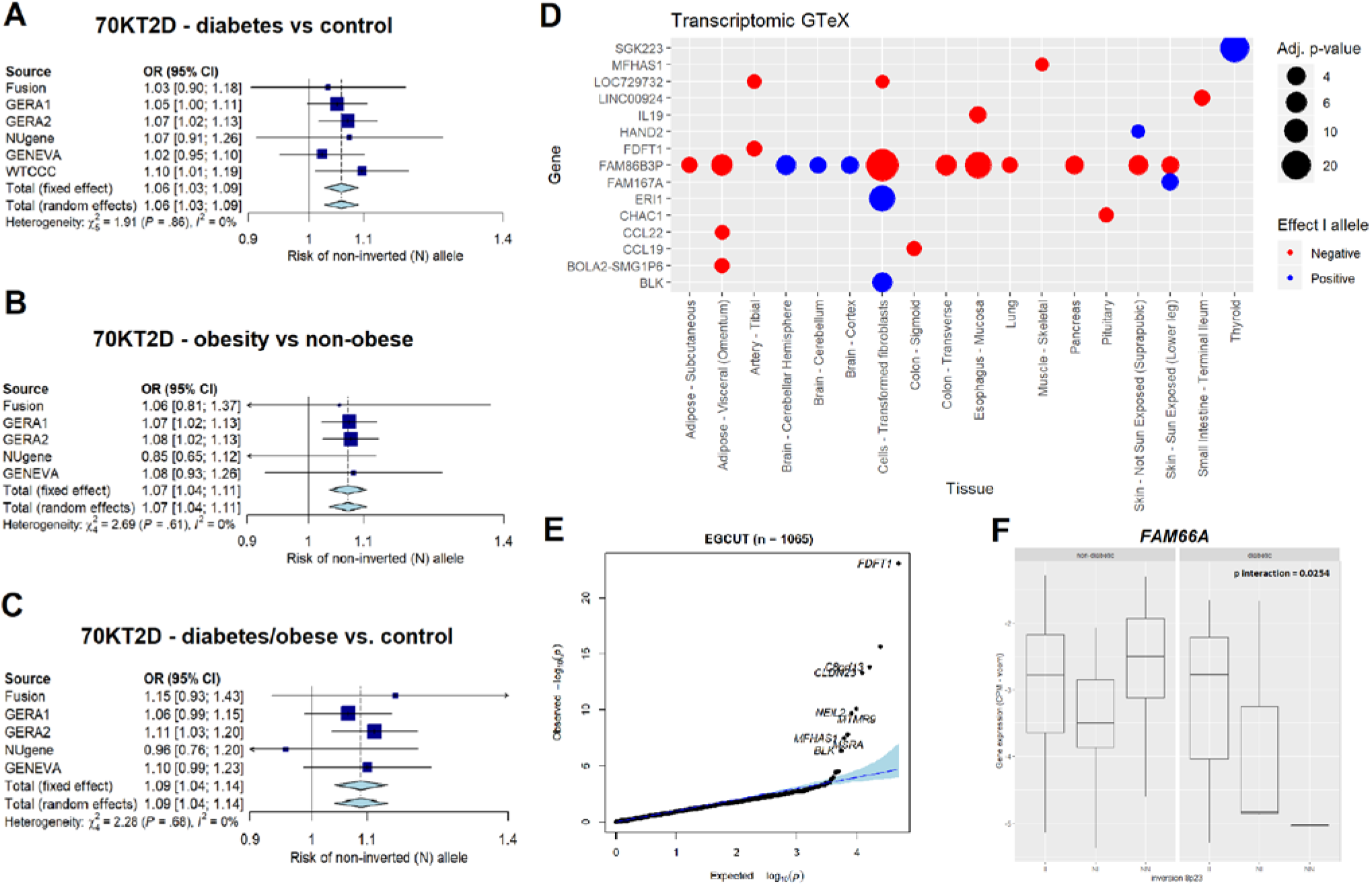
Validation of positive associations between the inversion 8p23.1 with diabetes, obesity and their co-occurrence in the 70KT2D dataset and transcriptional allelic effects in samples from EGCUT Biobank and GTEx tissues. **A**: Meta-analysis of datasets belonging to 70KT2D for the association of inversion 8p23.1 with diabetes. **B**: Meta-analysis of datasets belonging to 70KT2D for the association of inversion 8p23.1 with obesity. **C**: Meta-analysis of datasets belonging to 70KT2D for the association of inversion 8p23.1 with obese and diabetic individuals. **D:** Differential expressed genes at inversion genotypes (at 5% FDR) in different tissues from GTEx. **E:** Differentially expressed genes at inversion genotypes (at 5% FDR) in blood samples from EGCUT Biobank. **F:** *FAM66A* gene expression interaction between diabetic status and inversion 8p23.1 in pancreatic islets samples (p=0.0254).

Some interesting new associations in the discovery sample included those of inversion 17q21.31 with HDL, waist circumference, waist-hip ratio and systolic and diastolic blood pressure (**Figure 1**). Interestingly, this inversion also showed a significant role in behavioral traits such as mood swing, depression and bipolar disorder, which would need further validation. While we also found significant association of the inversion 6p21.33 with asthma (OR=1.02, p=0.0215) and different respiratory capacity traits (FEV1, p= 3.4×10^−9^ and FVC, p=3.2×10^−9^) the association with asthma was not replicated in the GERA study. The inversion 7q11.22 was associated with different morphometric traits (BMI, waist circumference and waist-to-hip ratio) and will require further validation studies.

### Association of inversions at 8p23.1, 16p11.2 with the co-occurrence of diseases is significant at genome-wide level while single SNPs are not

Remarkably, the N-allele of the inversion 8p23.1 was significantly associated with the co-occurrence of obesity with diabetes (OR=1.08, p=3.1×10^−7^), hypertension (OR=1.07, p=1.7×10^−16^), asthma (OR=1.08, p=3.0×10^−11^) and depression (OR=1.04, p=1.4×10^−3^). These results were validated in the GERA and 70KT2D (**Table 2**). For obesity/diabetes we observe an OR=1.17 (p=1.4×10^−13^) (**Figure 2C**) and none of the SNPs located within the inverted region were significantly associated at a genome-wide level (minimum p = 3.8×10^−5^) (**Supplementary Figure S4A**). Finally, we also found a significant association of the N-allele of inversion 11q13.2 with the co-occurrence of obesity with diabetes (OR=1.05, p=0.0011) and hypertension (OR=1.03, p=2.9×10^−5^) (**Figure 1**) which was not validated in the GERA study.

Some relevant findings have also been obtained in the study of inversion 16p11.2. These include new significant associations between the inversion and the co-occurrence of obesity with several diseases (**Figure 1).** The co-occurrence with diabetes at UKB (OR=1.06, p=7.5×10^−5^) was independently replicated in the 70KT2D study (OR=1.13, p=1.2×10^−8^) were none of the SNPs located within the inverted region were significantly associated at a genome-wide level (minimum p: 0.0214) (**Supplementary Figure S4B**). In addition, the significant co-occurrence with hypertension observed in the UKB study (OR=1.06, 2.7×10^−14^) was validated in the GERA study (OR=1.05, p=0.0357) further confirming the robustness of these findings (see **Table 2** reporting the effect of the risk allele N).

### Regulatory region and gene disruption are the mechanisms underlying the effect of inversions on diabetes

To investigate the possible mechanisms underlying the shared genetic influences of the inversions with obesity and its co-morbidities, we analyzed the transcriptional effects of the 21 inversions on different tissues from the GTEx project (see **Methods**). As a result of these analyses, we found that inversion 8p23.1 modulated the transcription in brain, pancreas and adipose tissue of *FAM86B3P* gene, as well as *SGK223, MFHAS1, IL19, HAND2, FDFT1, FAM167A, ERI1, CHAC1, CCL22, CCL19 and BLK* in other tissues (**Figure 2D**). Genes *FDFT1, C8orf13, CLDN23, NEIL2*, *MTMR9*, *MFHSA1, MSRA, BLK* and were also differentially expressed in blood samples from the validation study we performed in the independent general population cohort belonging to EGCUT Biobank (**Figure 2E**). For the inversion 16p11.2 we found a total of 30 genes differentially expressed at 5% FDR level in blood, brain, pancreas or adipose tissue including *TUFM, SULT1A2, SPNS1, EIF3CL, FOXO1* among many others (**Supplementary File 2**). These results were also observed in the blood samples of the validation cohort from EGCUT Biobank (**Supplementary Figure S5**). The genes affected by the other inversions and the different tissues can be found in **Supplementary File 2**.

A more detailed analysis of gene expression has been conducted on a relevant tissue to support the association on diabetic/obese patients. We first genotyped the inversions and analyzed RNA sequencing in human pancreatic islets from 89 deceased donors (see **Methods**). This revealed a significant association between inversion 8p23.1 and the expression levels of *CLDN23* (p=1.3×10^−3^) and *ERI1* (p=0.0356). We observed a nominally significant interaction of inversion 8p23.1 with obese/diabetic status associated with the expression of lncRNA *FAM66A* (p= 0.0254), where individuals carrying the risk allele for obesity and diabetes also present *FAM66A* downregulation (**Figure 2F**). In addition, results with inversion at 16p11.2 also revealed a significant interaction between the inversion and obese/diabetic status for the expression of *NUPR1* (p= 0.0116) and *ATXN2L* (p= 0.0167) (**Supplementary Figure S6**).

We also investigated whether the positional effects of the inversions could be associated with diabetes (see **Methods**). **Figure 3A** shows the chromatin landscape of the region of the inversion 8p23.1 as well as the location of all genes having a significant alteration of expression, including those that are islet-specifically expressed. A cluster of islet-specific genes is located outside the rightmost boundary of the inversion but inside the inversion’s topologically associated domains (TAD). Therefore, it is likely that the regulatory regions of these genes lie across the inversion’s boundary, and thus their *cis*-regulatory SNPs being separated from their target genes by the right breakpoint of the inversion 8p23.1 in the case of genes *FAM66A* and *FAM66D* (**Figure 3A**). Similarly, the analysis of the inversion 16p11.2 also revealed four eQTLs in which the *cis*-regulatory SNPs were separated from their target genes by the inversion breakpoints: *TUFM*, *SULT1A1*, *EIF3C EIF3CL* (**Figure 3B**). It is also worth to notice that *EIF3CL* gene is disrupted by the inversion breakpoint providing a different mechanism of action for that gene (**Figure 3B**).

**Figure 3.**
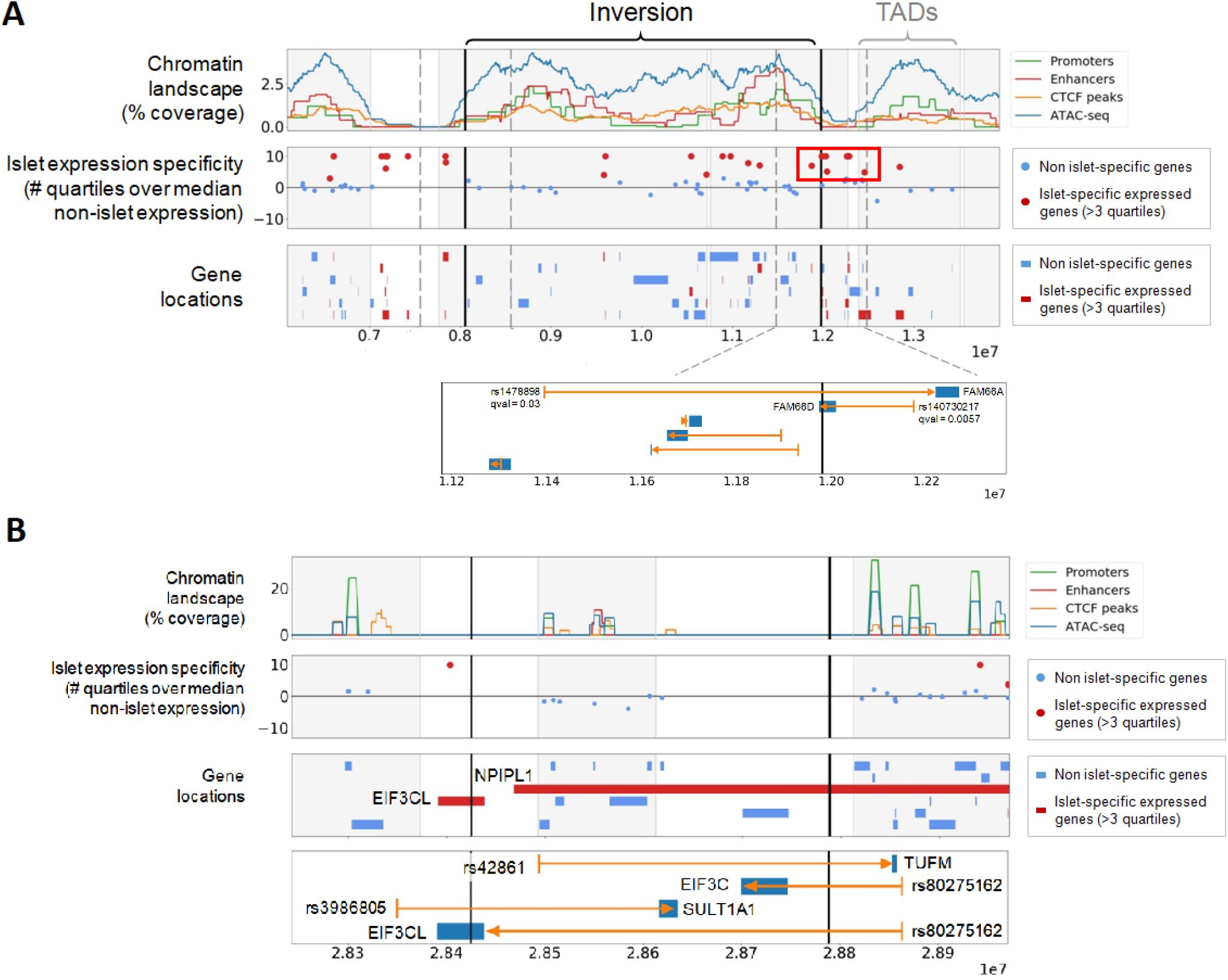
Mechanisms underlying the inversion association with diabetes. **Panel A** shows the islet specific expression of inversion 8p23.1 genes. We observed a cluster of islet-specific genes, mainly lncRNAs, next to the distal inversion breakpoint that could be separated from regulatory elements located inside the inverted region. The bottom panel depicts an eQTLs (rs1478898) of *FAM66A* gene disrupted by the inversion distal breakpoint (van de Bunt et al, 2015). *FAM66D* has its gene body split in two by the inversion, and would also have its promoter separated from its eQLT SNP (rs140730217) by the inversion. This could be the most likely causal candidate. **Panel B** shows the same information for the inversion 16p11.2. *TUFM* and *EIF3C* have their lead eQTL SNP separated by the inversion breakpoint. There is no evidence in the centiSNP database (Pique-Regi et al., 2016) for SNP rs42861 to be causal, suggesting that it should be in LD with the causal variant. This promoter region SNP is located in a segmental duplication block that is closer to *TUFM* in the inverted haplotypes. Therefore, positional changes made by the inversion can affect *TUFM* gene expression by separating the gene from regulatory sequences and subsequently increasing obesity risk.

### Obesity mediates the association of inversions with diabetes and hypertension

We first aimed to disentangle the shared genetic influence of the inversion 8p23.1 in obesity and diabetes. To this end, a Bayesian network analysis was performed on the discovery study (see **Methods**). Data indicated that the most likely model, based on the BIC, was for the sequence inv8p23.1 -> obesity -> diabetes, suggesting the mediatory effect of obesity on the association between the inversion and diabetes (**Figure 4A**). This was consistent with mediation analyses showing that 38.7% (CI95%: 25.2-59.0%) of the diabetes risk variance explained by the inversion 8p23.1 was mediated by obesity (p<10^−16^). Then, we also investigated whether inversion 8p23.1, 16p11.2 and 11q13.2 act jointly or not on obesity, diabetes and hypertension. The Bayesian network analysis including the three inversions in the model revealed that the inversions 8p23.1 and 16p11.2 independently associated with diabetes and hypertension being mediated by obesity (**Figure 4B**).

**Figure 4.**
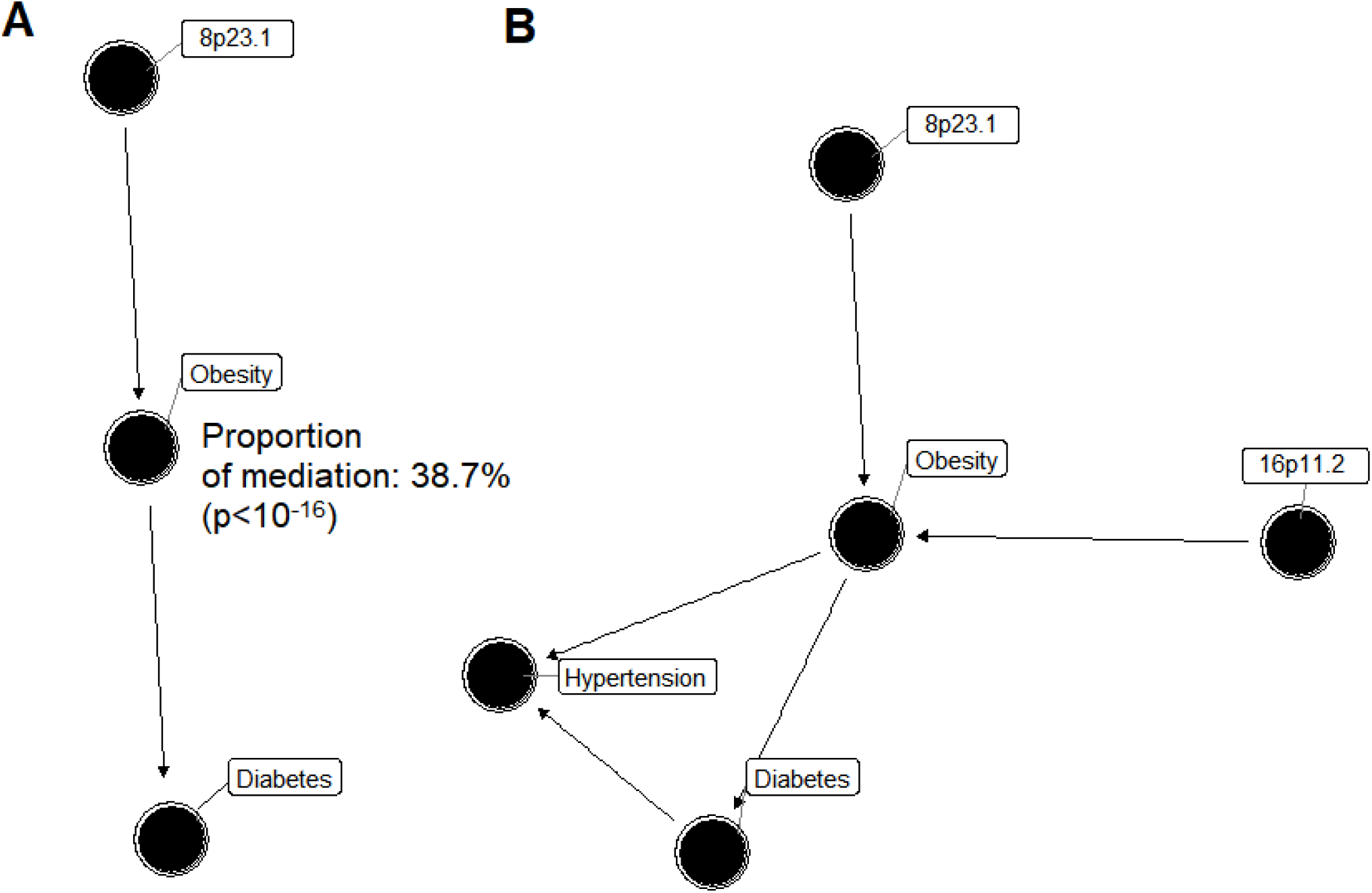
Mediation effect of obesity in the causal link between inversions and diabetes and hypertension. **Figure A** shows mediation analysis of obesity in the association between inversion 8p23.1 which is the Best Bayesian Network when analyzing these three variables. **Figure B** shows the Best Bayesian Network based on AIC obtained after including obesity, hypertension, diabetes and inversions 8p23.1, 16p11.2 and 11q13.2. Results are obtained from UKB data.

## DISCUSSION

Epidemiological studies largely support the co-occurrence of obesity with numerous traits and diseases such as diabetes, hypertension, asthma and psychiatric disorders among others^17,18^. The extent to which obesity is a cause, a consequence or shares common causes with these traits is subject of intense research^19–21^. Here, we show that at least two common polymorphic inversions at 8p23.1 and 16q11.2 offer a genetic substrate to some widely observed co-morbidities of obesity, such as those with diabetes, hypertension, asthma and depression. The analysis of UKB dataset validated the estimated inversion allele frequencies in European populations reported in our recent analyses^15^. The observed differences of some inversion allele frequencies among major populations could explain part of the existing geographic variability in disease incidence^22^. In particular, the reported cline of the inversion at 8p23.1 and 16p11.2 could capture a proportion of the observed North-South European differences in obesity^23^, diabetes and hypertension^24^ incidence.

The analysis of our discovery sample also confirmed previous reported associations of inversions with phenotypes, such as neuroticisms for the inversions 17q21.31 and 8p23.1^5^, obesity for inversion 8p23.1^3^, and the co-occurrence of asthma and obesity with the inversion 16p11.2^11^. In addition, we discovered and robustly validated new associations of the inversion 8p23.1 with diabetes and hypertension as well as the co-occurrences of obesity with diabetes, hypertension asthma and depression. These results suggest a relevant role of the inversion 8p23.1 in this metabolic syndrome^25^.

Our data suggest a causal path in which obesity mediates the observed association between inversions and several complex diseases. In particular, obesity mediates the independent effect of inversions at 8p23.1 and 16p11.2 on diabetes. Transcriptome analyses from general population has revealed candidate genes to mediate this effect, such as *BLK,* involved in pancreatic β-cell insulin metabolism whose rare mutations are associated with young age of onset diabetes^26^, or *FDFT1,* linked to C-reactive protein (CRP) and lipids levels^27^ and one of the strong candidates for obesity in gene expression networks derived from mouse intercrosses^28^. A more specific analysis of transcriptome and eQTLs on pancreatic islets leads to another interesting gene: *FAM66A*. *FAM66* is a multiple copy non-coding gene located in the flanking segmental duplications of the 8p23.1 inversion highly expressed in brain and with low-level expression in pancreas. Diabetic individuals carrying the N-allele have lower gene expression, while no differential expression across inversion genotypes is observed in control individuals. Therefore, allele-specific expression analysis of this gene shows clear differences in expression in pancreatic cells of already symptomatic diabetic patients. Remarkably, a copy-number gain variant including *FAM66* genes has been associated with increased risk of diabetes^29^. Positional analyses also pointed out *FAM66D* (8p23.1) as promising candidate since their gene body was split in two by the inversion breakpoint.

We have also shown that inversion at 16p11.2 affects the joint effect of obesity with diabetes and hypertension and that this effect is independent of the effect found for inversion 8p23.1. The functional consequences of this inversion were previously reported to be mediated by deregulation of *TUFM*, *SULT1A1*, *SULT1A2*, *SH2B1*, *APOB48R*, and *EIF3C* in blood^11^. Position transcriptional analysis in pancreatic islets revealed that *TUFM* and *EIF3C* have their lead eQTL SNPs separated in the inverted allele. Remarkably, the eQTL SNP rs42861 of *TUFM* does not seem to be causal in the centiSNP database^30^ suggesting that it is in LD with the causal variant. This SNP is located into the promoter region that is closer to the *TUFM* in the inverted haplotypes. This supports the hypothesis that the positional changes made by the inversion can affect *TUFM* gene expression and subsequently have an effect in obesity/diabetes increased risk. Positional analyses also pointed out *EIF3CL* since its gene body was split by the inversion breakpoint. This gene is an excellent candidate given that some of its isoforms were preferentially expressed in human pancreatic islets^31^.

The inversions at 8p23.1 and 16p11.2 also encompass the joint occurrence of obesity with behavioral traits, in particular with depression. The GTEx data analyses concede importance to *FAM86B3P* a pseudogene that has been associated with schizophrenia^32^ and neuroticism^33^, but also with type 2 diabetes^34^. These data further support our hypothesis that polymorphic inversions are strong candidates for the joint genetic susceptibility to co-occurring diseases by simultaneously affecting multiple genes. The observation that SNPs located in both inversion regions are not or weakly associated with the analyzed traits, while inversion haplotypes are associated even at genome-wide significant level for GWAS, and the strongest association found in people having more than one disease, also confirm that inversions are main contributors to the shared genetic susceptibility of co-occurring diseases. Functional analyses in the appropriate tissue in cases and controls, as the one we performed for obesity and diabetes, will shed light into the genes and mechanisms involved in behavioral or psychiatric traits.

Our hypothesis that inversions underly shared genetic susceptibility to common diseases is particularly supported by our findings in large inversions. These inversions encapsulated large numbers of genes and their associations with phenotypes were highly significant and could be replicated. Smaller inversions showed significant effects for numerous traits in the discovery study but only one result could be confirmed, namely the correlation of inversion at 11q13.2 with obesity and related traits and also with the co-occurrence of obesity, hypertension and diabetes. Similarly, this study opens the door to further inversion association studies for those traits not studied in this work. Additionally, the large number of significant genes associated with different tissues as well as the significant associations found for some traits also provides good candidate human diseases that are likely under the influence of inversions. These include, among others, Autism, Alzheimer and Parkinson disease.

In conclusion, our study is so far the largest association study regarding the number of inversions and human traits tested and represents a breakthrough for genomic association and comorbidity studies, in which polymorphic inversions were often previously disregarded. Our results underscore the role of some inversions as major genetic contribution to the joint susceptibility to common diseases. The results in obesity and diabetes provide, to our knowledge, the first example of a mechanism in which cis-regulatory SNPs are separated from their target genes by inversion breakpoints. These findings, set a new framework for future studies which is now accessible to the research community thanks to inversion genotyping tools such as our scoreInvHap method^15^.

## ONLINE METHODS

### Discovery Dataset

The UK Biobank (UKB) is a population-based cohort involving 500,000 individuals aged between 37 and 73 years, recruited across UK in the period 2006-2010. Further details on the quality control and genotyping are described in the study design^35^. Phenotypic information is recorded via questionnaires and interviews (e.g., demographics and health status) and SNP genotypes were generated from the Affymetrix Axiom UK Biobank and UKBiLEVE arrays. We based our study on 408,898 individuals from European descent and from whom inversion genotypes were called using SNP array data. Principal components were used in the analysis to control for population stratification.

### Replication Datasets

The GERA cohort (dbGaP Study Accession: phs000674.v1.p1 access granted to the authors) consists on individuals from Northern California Region (USA). Only individuals with reported race (variable phv00196837.v2.p2) equal to white were selected for the analyses. Individuals were genotyped with Affymetrix Axiom_KP_UCSF_EUR. After quality control of the inversion genotyping calling process a total of 53,782 individuals with information about sex, age, principal components for genetic ancestry and several diseases including obesity (9,439 cases), diabetes (6,529 cases), hypertension (27,009 cases), asthma (8,716 cases) and depression (6,924 cases) were used to replicate positive findings in the UKB cohort.

We further replicated the associations between inversions and type 2 diabetes and obesity in the 70KforT2D study (70KT2D)^36^. This study includes 5 datasets publicly available in dbGAP or EGA (NuGENE, FUSION, GENEVA, WTCCC and GERA). We used information about being diabetic or not as describe elsewhere^36^. The 5 datasets were used to replicate the significant findings in the UK Biobank data on diabetes. The WTCCC dataset was removed from the obesity and obesity/diabetes analysis since we did not have access to body mass index (BMI) information for that study. The GERA dataset was split in two (GERA1 and GERA2) to speed up the imputation and inversion calling procedure since it is a large dataset. After performing QC on inversion genotypes, a total of 67,299 individuals were used in the replication step (54,801 controls and 12,498 diabetic). Data was accessed from the portal cg.bsc.es/70kfort2d.

The obesity variable was created using body mass index (BMI) variable. Control individuals were considered those having BMI in the interval (18.5–24.9) and obese people were considered those having BMitalic>30.0. To replicate obesity associations, we excluded individuals with diabetes. As a result, a total of 34,316 individuals (23,818 controls and 10,498 obese) were used for that purpose. The co-occurrence of obesity and diabetes was studied by comparing individuals with no obesity and no diabetes as the reference category with individuals being obese and diabetic simultaneously. This ended up with a total of 23,818 control and 5,715 obese/diabetic individuals.

Next, we further describe the studies included in the 70KT2D dataset along with their accession numbers. These studies were granted for access to the co-authors.

#### Northwestern NUgene Project: Type 2 Diabetes (NUGENE)

NUgene project (dbGaP Study Accession: phs000237.v1.p1) contains data from patients from the Northwestern University Medical Center (USA). For this study, T2D cases were included if they had been diagnosed of Type 2 Diabetes, they took drugs to treat Type 2 Diabetes or they presented abnormal diabetes-related blood measures. Controls were included if they had not been diagnosed of Type 2 Diabetes, they did not take drugs to treat Type 2 Diabetes, they presented normal diabetes-related blood measures and they did not have any family history of diabetes (either Type 1 or Type 2). In both groups, patients with Type 1 Diabetes were excluded. These individuals were genotyped with Illumina Human1M-Duov3_B.

#### The Finland-United States Investigation of NIDDM Genetics - GWAS Study (FUSION)

FUSION study (dbGaP Study Accession: phs000100.v4.p1) aims to investigate the association between genetics and Type 2 Diabetes in Finish families. For this study, T2D cases were included if they had been diagnosed of Type 2 Diabetes, they took drugs to treat Type 2 Diabetes or they presented abnormal diabetes-related blood measures. Controls were included if they presented normal diabetes-related blood measures and were frequency matched to the cases by age, sex and birth province. In both groups, patients with Type 1 Diabetes familial history were excluded. These individuals were genotyped with Illumina HumanHap300v1.1.

#### GENEVA Genes and Environment Initiatives in Type 2 Diabetes (Nurses’ Health Study/Health Professionals Follow-up Study)

GENEVA project (dbGaP Study Accession: phs000091.v2.p1) is a nested case-control study from two USA cohorts: the Nurses’ Health Study (NHS) and the Health Professionals Follow-up Study (HPFS). These individuals were genotyped with Affymetrix AFFY_6.0.

#### Resource for Genetic Epidemiology Research on Aging (GERA)

GERA cohort (dbGaP Study Accession: phs000674.v1.p1) consists of individuals from Northern California Region (USA). These individuals were genotyped with Affymetrix Axiom_KP_UCSF_EUR.

### Geographical variation in Europe

POPRES project (dbGaP Study Accesion: phs000145.v4.p2 access granted to the authors) was used to estimate inversion frequencies in European countries and regions. This project aimed to facilitate exploratory genetic research by assembling a DNA resource from a large number of subjects participating in multiple studies throughout the world. We selected European individuals (variable phv00173964.v2.p2) leading a total of 3,071 samples. A geographic label (North, Center, South) was assigned to each individual using information of variable phv00066613.v2.p2.

### Transcriptomic analyses

#### GTEx Analysis

We associated the 21 chromosomal inversions to changes in gene expression in GTEx project. We determined inversion genotypes on the GTEx v7 genotype calls from dbGAP (dbGaP Study Accession: phs000424.v7.p2 accession granted to the authors). We only included samples classified as European with a confidence higher than 90% by peddy^37^. Inv3_003 was discarded as the calling was not confident. Gene expression counts from RNA-seq data were downloaded using recount2^38^. We computed the association between gene expression and inversions using voom^39^ and limma^40^. The linear model included the inversion coded as additive (0: NN, 1: NI, 2: II) and the same covariates than GTEx (first three genome-wide PCA components, sex and covariates from PEER). In each tissue, we selected those features having more than 10 counts in at least 10% of the samples. We corrected the association results per tissue for multiple comparison by using a false discovery rate (FDR) adjusted p-value per tissue.

#### EGCUT Biobank

Estonian Gene Expression Cohort (EGCUT, www.biobank.ee) was used to replicate positive transcriptomic results found in GTEx. The cohort is composed of 1,048 randomly selected samples (mean age 37+/-16.6 years; 50% females) from the cohort of 53,000 samples in the Estonian Genome Center Biobank, University of Tartu. Whole blood RNA sample was collected with Tempus Blood RNA Tubes (Life Technologies, NY, USA), RNA was extracted using Tempus Spin RNA Isolation Kit (Life Technologies, NY, USA) and quality was measured by NanoDrop 1000 Spectrophotometer (Thermo Fisher Scientific, DE, USA) and Agilent 2100 Bioanalyzer (Agilent Technologies, CA, USA) Whole-Genome gene-expression levels were obtained by Illumina HT12v3 arrays (Illumina Inc, San Diego, US) according manufactures protocols. Low quality samples were excluded. All probes with primer polymorphisms were discarded, leaving 34,282 probes. Raw gene expression data was Log-Quantile normalized using MixupMapper software. DNA was genotyped with Human370CNV array (Illumina Inc., San Diego, US).

#### Pancreatic Islets

We analyzed the transcriptomic effect of inversions 8p23.1 and 16q11.2 on 118 pancreatic human islet samples using RNA-sequencing counts and high-density genotyping data^41^. DNA genotype data (EGA accession number EGAS00001001261) was used to call inversion genotypes using *scoreInvHap*, then the association between gene expression and inversions was assessed using voom^39^ and limma^40^.

#### Positional analyses

For the positional analysis, several annotations were gathered from the following sources: TAD boundaries from the Human ES Cell (H1) topologial domains^42^; promoters, enhancers, CTCF-peaks and ATAC-seq open-chromatin regions from the human islet regulome annotation^43^; islet-specificity scores were calculated using the gene expression data from^31^; eQTL snp-gene associations from^41,44^. The chromatin landscape coverage percentage was calculated using a sliding window of 500kb and 1Mb for inversions 8p23.1 and 16p11.2 respectively, using steps of 1% of the window size, and calculating the percentage of covered nucleotides by significant signal in each of the categories. For the islet-specific expression analysis, we calculated the non-islet median expression level and difference between the 75 and 50 quartiles, and considered as islet-specific any gene that was expressed in islet >3 quartiles over the median of non-islet expression. Visualization was done in python3 using the matplotlib graphics library.

### Statistical methods

#### SNP imputation and inversion calling

SNP microarray data was imputed with *imputeInversion* pipeline prior to inversion calling^45^. This pipeline was designed to impute only those SNPs inside the inversion region or closer than 500 Kb to the inversion breakpoints an step that is recommended before performing inversion calling. *imputeInversion* uses shapeITv2.r904 to phase^46^, Minimac3^47^ to impute and 1000 Genomes as reference haplotypes. Variants with an imputation R2 < 0.3 were discarded. Genotype probabilities were used to call inversions using *scoreInvHap*^15^ which is available at Bioconductor. *scoreInvHap* computes a similarity score between an individual’s alleles and the reference alleles in each chromosomal status. We used the development version of *scoreInvHap*, which includes references for 21 inversions. These methods were used to perform inversion calling of discovery and replication studies as well as individuals from POPRES.

#### Inversion frequencies

Inversion frequencies were estimated in UKB and POPRES studies using SNPassoc package^48^. A trend test implemented in the R function *prop.trend.test* was used to assess whether inversion frequencies in European regions from POPRES (North, Center, South) had a significant cline. Principal component analysis was used to visualize inversion frequencies across European regions of POPRES dataset.

#### Obesity and obesity co-occurrence traits

Obesity trait was created using body mass index (BMI) information. First, BMI was categorized in 5 categories using World Health Organization (WHO) classification (http://www.euro.who.int/en/health-topics/disease-prevention/nutrition/a-healthy-lifestyle/body-mass-index-bmi) which considers the following categories: underweight (BMI below 18.5), normal weight (BMI between 18.5 and 25), pre-obesity (BMI between 25 and 29.9), obesity class I (BMI between 30 and 34.9) and obesity class II and III (BMI above 35). Obesity was considered as obesity class I, II and III and was compared with normal weight category. The analysis of obesity co-occurrence with diabetes, hypertension, asthma, depression and neuroticism was performed by comparing individuals with normal weight and no presence of the disease with individuals being obese and having the disease of interest.

#### Inversion association analyses

Each inversion was independently associated with all the traits by using generalized linear models implemented in SNPassoc package^48^. The models were adjusted for gender, age and the first four principal components obtained from GWAS data in order to control for population genetic differences. The inversions were analyzed using an additive model. Multiple comparison problem was considered by correcting for the total number of inversions analyzed.

#### Causal inference

Mediation analysis using *mediation* R package^49^ was used to evaluate whether inversion 8p23.1 mediates the association between obesity and diabetes. Additive Bayesian network models using *abn* R package^50^ was used to determine optimal Bayesian network models to identify statistical dependencies between inversions 8p23.1, 16p11.2 and 11q13.2 and obesity, diabetes and hypertension. The most probable network structure was estimated using exact order-based approach as implemented in the *mostprobable* function of *abn* package.

## ACKNOWLEDGEMENTS

This research has received funding from Ministerio de Ciencia, Innovación y Universidades (MICIU), Agencia Estatal de Investigación (AEI) and Fondo Europeo de Desarrollo Regional, UE (RTI2018-100789-B-I00); and the Catalan Government [SGR2017/801 and #016FI_B 00272 to CR-A]. JG is funded by the European Commission (H2020-ERC-2014-CoG-647900) and the MINECO/AEI/FEDER, EU (BFU2017-82937-P). LAPJ lab was funded by the Spanish Ministry of Science and Innovation (ISCIII-FEDER P13/02481), the Catalan Department of Economy and Knowledge (SGR2014/1468, SGR2017/1974 and ICREA Acadèmia), and also acknowledges support from the Spanish Ministry of Economy and Competiveness “Programa de Excelencia María de Maeztu” (MDM-2014-0370). This research has been conducted using the UK Biobank Resource under Application Number 43983. The Genotype-Tissue Expression (GTEx) Project was supported by the Common Fund of the Office of the Director of the National Institutes of Health, and by NCI, NHGRI, NHLBI, NIDA, NIMH, and NINDS.

## Contributions

JRG designed and oversaw the study and helped with data analyses. CR-A, AC, ML and IT performed quality control and inversion calling of UK Biobank, GERA and 70KT2D datasets. MG-M and JMM downloaded and pre-processed GERA data. TE analysed transcriptomic EGCUT data. IM, LA and DT downloaded and pre-processed 70KT2D data and performed transcriptomic analyses of pancreatic human islet samples. IM carried out positional analyses of inversions. JRG, JG and LAP-J wrote the manuscript with feedback from all authors.

## Data availability

The data used in this work were obtained from publicly available datasets that are accessible through public repositories: UKB study, dbGaP, EGA, GTeX and GEO. The inversion calling of UKB samples will be available through their platform. The inversion calling for the other samples are available upon request. The complete transcriptomic summary statistics of the 21 inversions are also available upon request. We plan to make publicly available this information in a Shiny app before the paper is accepted.

## Code availability

We made use of publicly available software and tools. An R Markdown document used to generate results that are reported in this paper is available upon request.

## References

1. Collaborators, T. G. 2015 O. Health Effects of Overweight and Obesity in 195 Countries over 25 Years. N. Engl. J. Med. 377, 13–27 (2017).

2. Dixon, J. B. The effect of obesity on health outcomes. Mol. Cell. Endocrinol. 316, 104–8 (2010).

3. Cáceres, A. & González, J. R. Following the footprints of polymorphic inversions on SNP data: from detection to association tests. Nucleic Acids Res. 1–11 (2015). doi:10.1093/nar/gkv073

4. Gutiérrez Arumi, A. Ancestral genomic submicroscopic inversions of human genome and their relation with multifactorial human diseases. Univ. Pompeu Fabra (2015).

5. Okbay, A. et al. Genetic variants associated with subjective well-being, depressive symptoms, and neuroticism identified through genome-wide analyses. Nat. Genet. 48, 624–33 (2016).

6. Karlsson Linnér, R. et al. Genome-wide association analyses of risk tolerance and risky behaviors in over 1 million individuals identify hundreds of loci and shared genetic influences. Nat. Genet. 51, 245–257 (2019).

7. Laws, S. M. et al. Fine mapping of the MAPT locus using quantitative trait analysis identifies possible causal variants in Alzheimer’s disease. Mol. Psychiatry 12, 510–517 (2007).

8. Zabetian, C. P. et al. Association analysis of MAPT H1 haplotype and subhaplotypes in Parkinson’s disease. Ann. Neurol. 62, 137–144 (2007).

9. Pilbrow, A. P. et al. Cardiac CRFR1 Expression Is Elevated in Human Heart Failure and Modulated by Genetic Variation and Alternative Splicing. Endocrinology 157, 4865–4874 (2016).

10. Ikram, M. A. et al. Common variants at 6q22 and 17q21 are associated with intracranial volume. Nat. Genet. 44, 539–544 (2012).

11. González, J. R. et al. A common 16p11.2 inversion underlies the joint susceptibility to asthma and obesity. Am. J. Hum. Genet. 94, (2014).

12. de Jong, S. et al. Common inversion polymorphism at 17q21.31 affects expression of multiple genes in tissue-specific manner. BMC Genomics 13, 458 (2012).

13. Chaisson, M. J. P. et al. Multi-platform discovery of haplotype-resolved structural variation in human genomes. Nat. Commun. 10, 1784 (2019).

14. Giner-Delgado, C. et al. Evolutionary and functional impact of common polymorphic inversions in the human genome. Nat. Commun. 10, 4222 (2019).

15. Ruiz-Arenas, C. et al. scoreInvHap: Inversion genotyping for genome-wide association studies. PLOS Genet. 15, e1008203 (2019).

16. Pickrell, J. K. et al. Detection and interpretation of shared genetic influences on 42 human traits. Nat. Genet. 48, 709–717 (2016).

17. Banks, J., Marmot, M., Oldfield, Z. & Smith, J. P. Disease and Disadvantage in the United States and in England. JAMA 295, 2037 (2006).

18. Stunkard, A. J., Faith, M. S. & Allison, K. C. Depression and obesity. Biol. Psychiatry 54, 330–7 (2003).

19. Martins-Silva, T. et al. Assessing causality in the association between attention-deficit/hyperactivity disorder and obesity: a Mendelian randomization study. Int. J. Obes. 1 (2019). doi:10.1038/s41366-019-0346-8

20. Xu, S., Gilliland, F. D. & Conti, D. V. Elucidation of causal direction between asthma and obesity: a bi-directional Mendelian randomization study. Int. J. Epidemiol. (2019). doi:10.1093/ije/dyz070

21. Millard, L. A. C., Davies, N. M., Tilling, K., Gaunt, T. R. & Davey Smith, G. Searching for the causal effects of body mass index in over 300 000 participants in UK Biobank, using Mendelian randomization. PLOS Genet. 15, e1007951 (2019).

22. Puig, M., Casillas, S., Villatoro, S. & Cáceres, M. Human inversions and their functional consequences. Brief. Funct. Genomics 14, 369–379 (2015).

23. Berghöfer, A. et al. Obesity prevalence from a European perspective: a systematic review. BMC Public Health 8, 200 (2008).

24. Wolf-Maier, K. et al. Hypertension Prevalence and Blood Pressure Levels in 6 European Countries, Canada, and the United States. JAMA 289, 2363 (2003).

25. Povel, C. M., Boer, J. M. A., Reiling, E. & Feskens, E. J. M. Genetic variants and the metabolic syndrome: a systematic review. Obes. Rev. 12, 952–967 (2011).

26. Borowiec, M. et al. Mutations at the BLK locus linked to maturity onset diabetes of the young and -cell dysfunction. Proc. Natl. Acad. Sci. 106, 14460–14465 (2009).

27. Ligthart, S. et al. Bivariate genome-wide association study identifies novel pleiotropic loci for lipids and inflammation. BMC Genomics 17, 443 (2016).

28. Logsdon, B. A., Hoffman, G. E. & Mezey, J. G. Mouse obesity network reconstruction with a variational Bayes algorithm to employ aggressive false positive control. BMC Bioinformatics 13, 53 (2012).

29. Bailey, J. N. C. et al. The Role of Copy Number Variation in African Americans with Type 2 Diabetes-Associated End Stage Renal Disease. J. Mol. Genet. Med. 7, 61 (2013).

30. Moyerbrailean, G. A. et al. Which Genetics Variants in DNase-Seq Footprints Are More Likely to Alter Binding? PLOS Genet. 12, e1005875 (2016).

31. Miguel-Escalada, I. et al. Human pancreatic islet three-dimensional chromatin architecture provides insights into the genetics of type 2 diabetes. Nat. Genet. 51, 1137–1148 (2019).

32. Li, Z. et al. Genome-wide association analysis identifies 30 new susceptibility loci for schizophrenia. Nat. Genet. 49, 1576–1583 (2017).

33. Luciano, M. et al. Association analysis in over 329,000 individuals identifies 116 independent variants influencing neuroticism. Nat. Genet. 50, 6–11 (2018).

34. Xue, A. et al. Genome-wide association analyses identify 143 risk variants and putative regulatory mechanisms for type 2 diabetes. Nat. Commun. 9, 2941 (2018).

35. Sudlow, C. et al. UK Biobank: An Open Access Resource for Identifying the Causes of a Wide Range of Complex Diseases of Middle and Old Age. PLOS Med. 12, e1001779 (2015).

36. Bonàs-Guarch, S. et al. Re-analysis of public genetic data reveals a rare X-chromosomal variant associated with type 2 diabetes. Nat. Commun. 9, 321 (2018).

37. Pedersen, B. S. & Quinlan, A. R. Who’s Who? Detecting and Resolving Sample Anomalies in Human DNA Sequencing Studies with Peddy. Am. J. Hum. Genet. 100, 406–413 (2017).

38. Collado-Torres, L. et al. Reproducible RNA-seq analysis using recount2. Nat. Biotechnol. 35, 319–321 (2017).

39. Law, C. W., Chen, Y., Shi, W. & Smyth, G. K. voom: Precision weights unlock linear model analysis tools for RNA-seq read counts. Genome Biol. 15, R29 (2014).

40. Ritchie, M. E. et al. limma powers differential expression analyses for RNA-sequencing and microarray studies. Nucleic Acids Res. 43, e47 (2015).

41. van de Bunt, M. et al. Transcript Expression Data from Human Islets Links Regulatory Signals from Genome-Wide Association Studies for Type 2 Diabetes and Glycemic Traits to Their Downstream Effectors. PLoS Genet. 11, e1005694 (2015).

42. Dixon, J. R. et al. Topological domains in mammalian genomes identified by analysis of chromatin interactions. Nature 485, 376–380 (2012).

43. Pasquali, L. et al. Pancreatic islet enhancer clusters enriched in type 2 diabetes risk–associated variants. Nat. Genet. 46, 136 (2014).

44. Fadista, J. et al. Global genomic and transcriptomic analysis of human pancreatic islets reveals novel genes influencing glucose metabolism. Proc. Natl. Acad. Sci. U. S. A. 111, 13924–9 (2014).

45. Tolosana, I., Ruiz-Arenas, C. & González, J. R. imputeInversion. (2018).

46. Delaneau, O., Zagury, J.-F. & Marchini, J. Improved whole-chromosome phasing for disease and population genetic studies. Nat. Methods 10, 5–6 (2013).

47. Das, S. et al. Next-generation genotype imputation service and methods. Nat. Genet. 48, 1284–1287 (2016).

48. González, J. R. et al. SNPassoc: an R package to perform whole genome association studies. Bioinformatics 23, 644–5 (2007).

49. Tingley, D., Yamamoto, T., Hirose, K., Keele, L. & Imai, K. **mediation**: *R* Package for Causal Mediation Analysis. J. Stat. Softw. 59, 1–38 (2014).

50. Lewis, F. I. & Ward, M. P. Improving epidemiologic data analyses through multivariate regression modelling. Emerg. Themes Epidemiol. 10, 4 (2013).

