## Supplementary Tables and Figures for "Polymorphic inversions underlie the shared genetic susceptibility to prevalent common diseases"

*Supplementary Material*

### Supplementary Tables

**Table S1**. **Descriptive data from UKB (N=408,898).** First column shows the observed frequency and percentage for categorical variables and median and standard deviation for continuous variables. The second column depicts the effective number of individuals used at each association analysis with the inversions. Obesity was considered as obesity class I, II and III using World Health Organization (WHO) classification. Individuals with normal weight and no presence of disease co-occurrence were used as the refence individuals for the disease/obesity co-occurrence analysis (see Methods).

| | | **[Effective samples]** ***N*** |
|---|---|---|
| **Obesity**: | | 258,950 |
| No | 133,372 (51.5%) | |
| Yes | 125,578 (48.5%) | |
| **BMI** | 27.4 (4.76) | 407,602 |
| **BMIcat**: | | 407,602 |
| Underweight (0,18.5) | 1,163 (0.29%) | |
| Normal weight [18.5-25) | 133,372 (32.7%) | |
| Pre-obesity [25-29.9) | 147,489 (36.2%) | |
| Obesity class I [30, 34.9) | 115,714 (28.4%) | |
| Obesity class II and III [35, Inf) | 9,864 (2.42%) | |
| **Waist circumference** | 90.3 (13.5) | 408,224 |
| **Hip** | 103 (9.15) | 408,179 |
| **Waist-hip ratio** | 0.87 (0.09) | 408,141 |
| **Diabetes**: | | 408,898 |
| No | 391,771 (95.8%) | |
| Yes | 17,127 (4.19%) | |

| | | [Effective samples] *N* |
|---|---|---|
| **Fasting Glucose** | 5.12 (1.21) | 356,598 |
| **HDL** | 1.45 (0.38) | 356,842 |
| **LDL** | 3.57 (0.87) | 389,160 |
| **Triglycerides** | 1.76 (1.02) | 389,569 |
| **Total Cholesterol** | 5.71 (1.15) | 389,877 |
| **Stroke**: | | 408,898 |
| No | 402,308 (98.4%) | |
| Yes | 6590 (1.61%) | |
| **Hypertension**: | | 408,898 |
| No | 298,288 (72.9%) | |
| Yes | 110,610 (27.1%) | |
| **Systolic blood pressure** | 140 (19.7) | 381,982 |
| **Diastolic blood pressure** | 82.3 (10.7) | 381,990 |
| **Asthma**: | | 408,898 |
| No | 360,131 (88.1%) | |
| Yes | 48,767 (11.9%) | |
| **COPD**: | | 408,898 |
| No | 406,809 (99.5%) | |
| Yes | 2,089 (0.51%) | |
| **fev1** | 2.84 (0.79) | 372,730 |
| **fvc** | 3.75 (1.05) | 372,730 |
| **fev1_fvc** | 0.42 (0.89) | 329,335 |
| **Depression**: | | 130,803 |
| No | 60,801 (46.5%) | |
| Yes | 70,002 (53.5%) | |
| **Bipolar disorder**: | | 96,832 |
| No | 69,980 (72.3%) | |
| Yes | 26,852 (27.7%) | |
| **Neuroticism**: | 4.11 (3.26) | 332,268 |
| **Risk Behaviour**: | | 395,109 |
| No | 295,107 (74.7%) | |
| Yes | 100,002 (25.3%) | |
| **Mood Swin**: | | 399,421 |
| No | 218,703 (54.8%) | |
| Yes | 180,718 (45.2%) | |
| **Diabetes/obesity**: | | 142,463 |
| No | 131,604 (92.4%) | |
| Yes | 10,859 (7.62%) | |

| | | [Effective samples] ***N*** |
|---|---|---|
| **Hypertension/obesity**: | | 164,388 |
| No | 112,949 (68.7%) | |
| Yes | 51,439 (31.3%) | |
| **Asthma/obesity**: | | 136,583 |
| No | 119,092 (87.2%) | |
| Yes | 17,491 (12.8%) | |
| **Depression/obesity**: | | 45,206 |
| No | 23,052 (51.0%) | |
| Yes | 22,154 (49.0%) | |
| **Neuroticism/obesity**: | | 10,4336 |
| No | 92,618 (88.8%) | |
| Yes | 11,718 (11.2%) | |

### Supplementary Figures

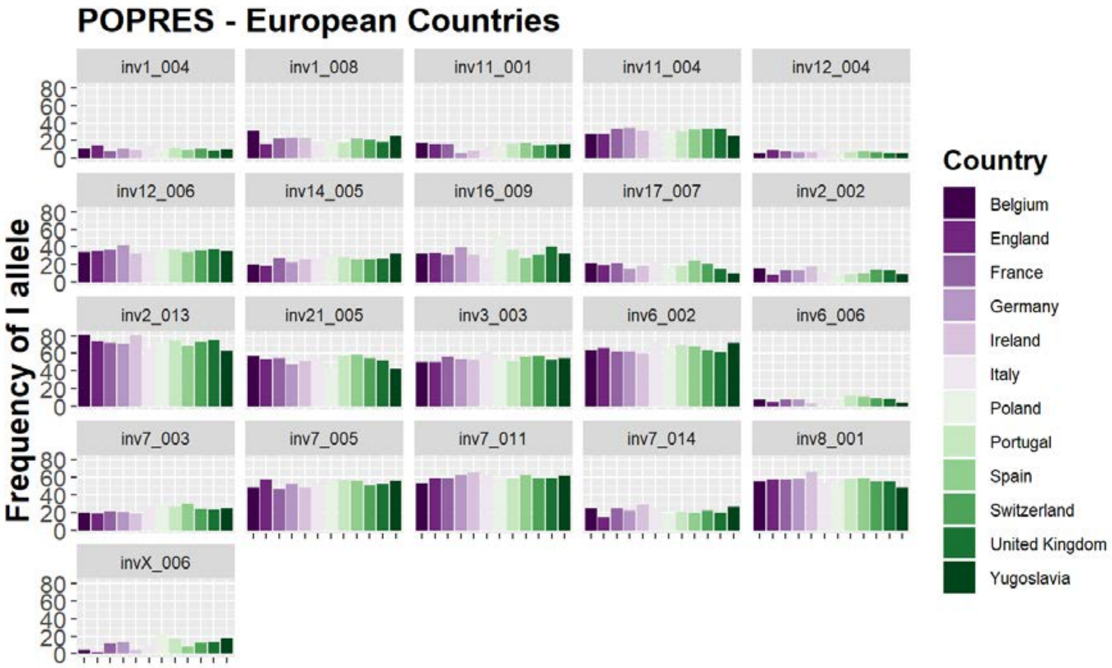


**Figure S1**. **Inversion frequencies at European countries.** Barplots shows the frequency of the I-allele for the 21 inversions analyzed in 12 countries from the POPRES study.

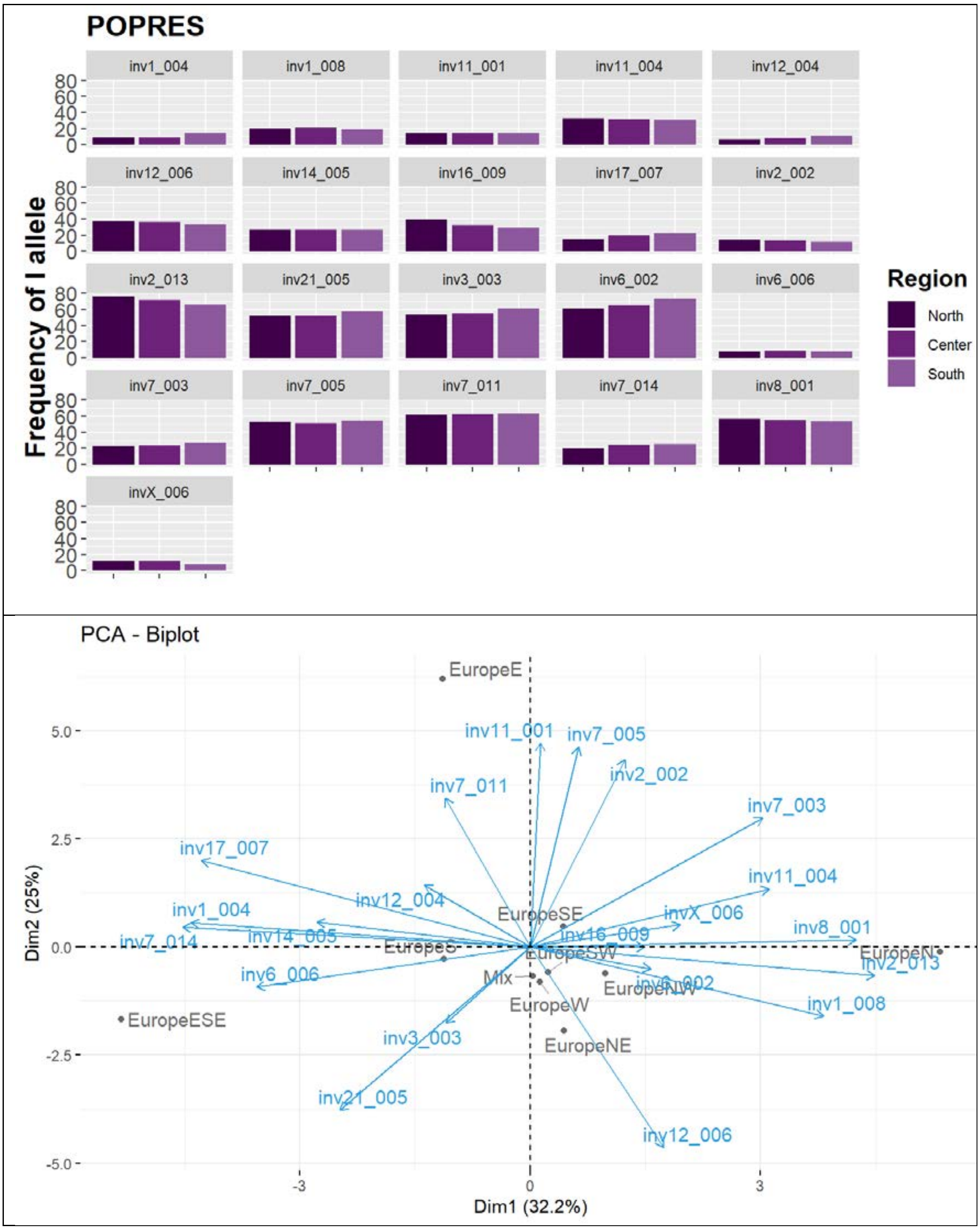


**Figure S2**. **Geographical distribution of inversion frequencies. Panel A** shows the frequency of I-allele of the 21 analyzed inversions by three European regions (North, Center and South) from POPRES study. **Panel B** shows the biplot obtained after performing a PCA of the I-allele frequencies by European regions from POPRES study. In both cases regions were obtained from variable GROUPING_PCA_LABEL2 computed at POPRES dataset.

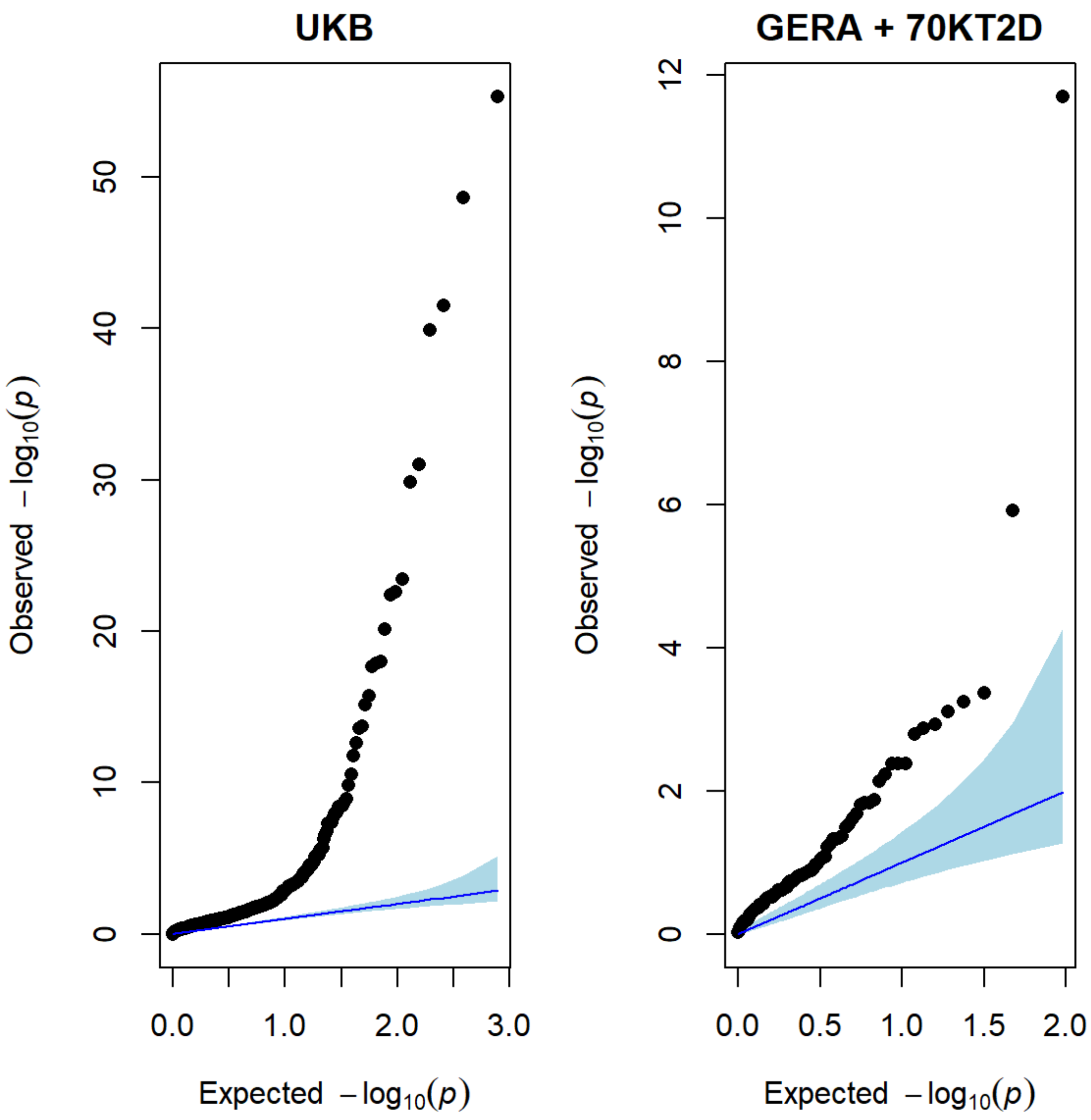


**Figure S3**. **Inflation in inversion association studies**. Q-Q plot of p-values obtained after assessing the association between 25 traits and diseases and 6 obesity co-occurrences with the 21 inversions from the discovery sample (**panel A**) and 5 diseases and 4 obesity co-occurrences with 21 inversions analyzed in GERA study (**panel B**).

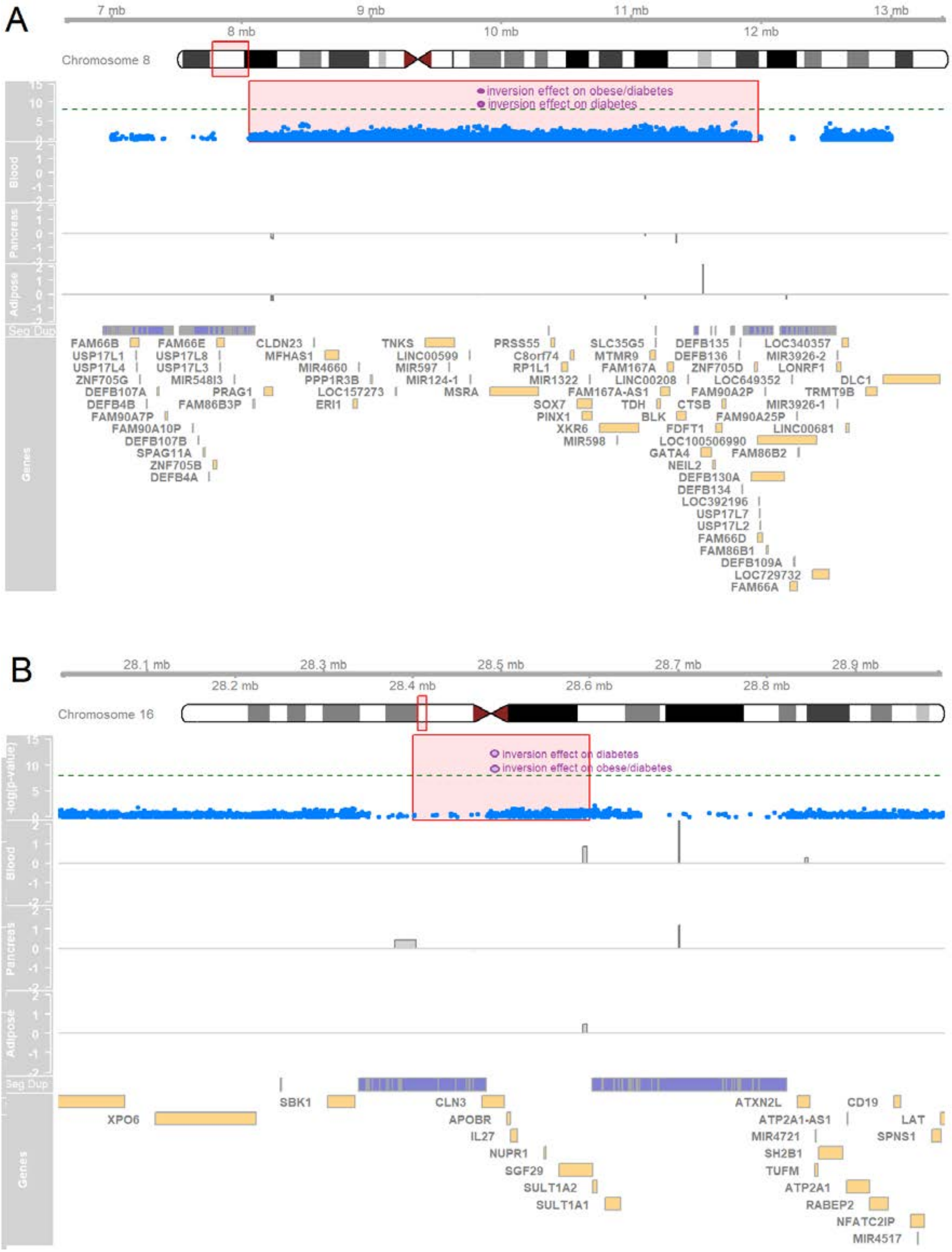


**Figure S4**. **GWAS results from 70KT2D study** (http://cg.bsc.es/70kfort2d/). The figures show the Manhattan plot corresponding to the -log10(p-values) of the association between the SNPs located in the inversions 8p23.1 (**panel A**) and 16p11.2 (**panel B**) and diabetes (blue dots). The significance for the inversions is in purple. The next two tracks show the significant genes associated with the inversions in GTEx in blood, pancreas and adipose tissue. Segmental duplications are then depicted in dark blue along with the annotated genes in the region that are showed in the last track.

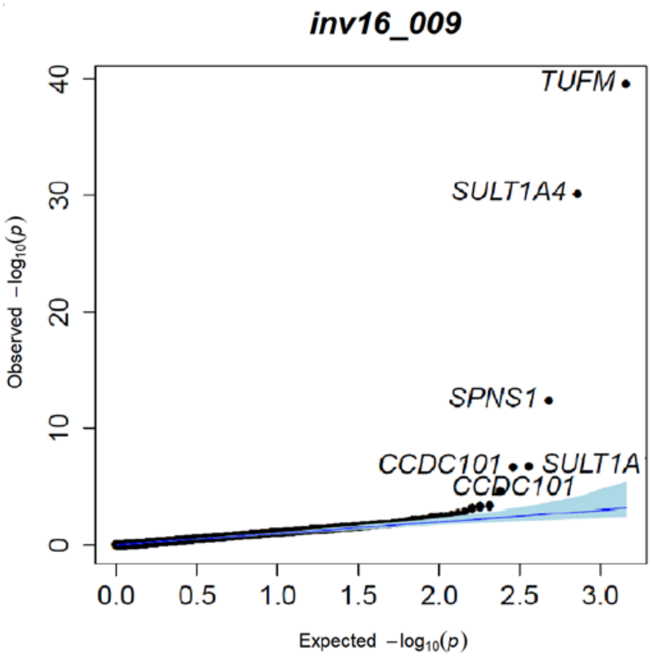


**Figure S5**. **Gene expression analysis in the EGCUT data set.** Transcriptomic analysis of blood samples from 1065 individuals belonging to EGCUT Estonian Biobank. Figure shows the significant associations for inversion 16p11.2 when analyzing the whole transcriptome.

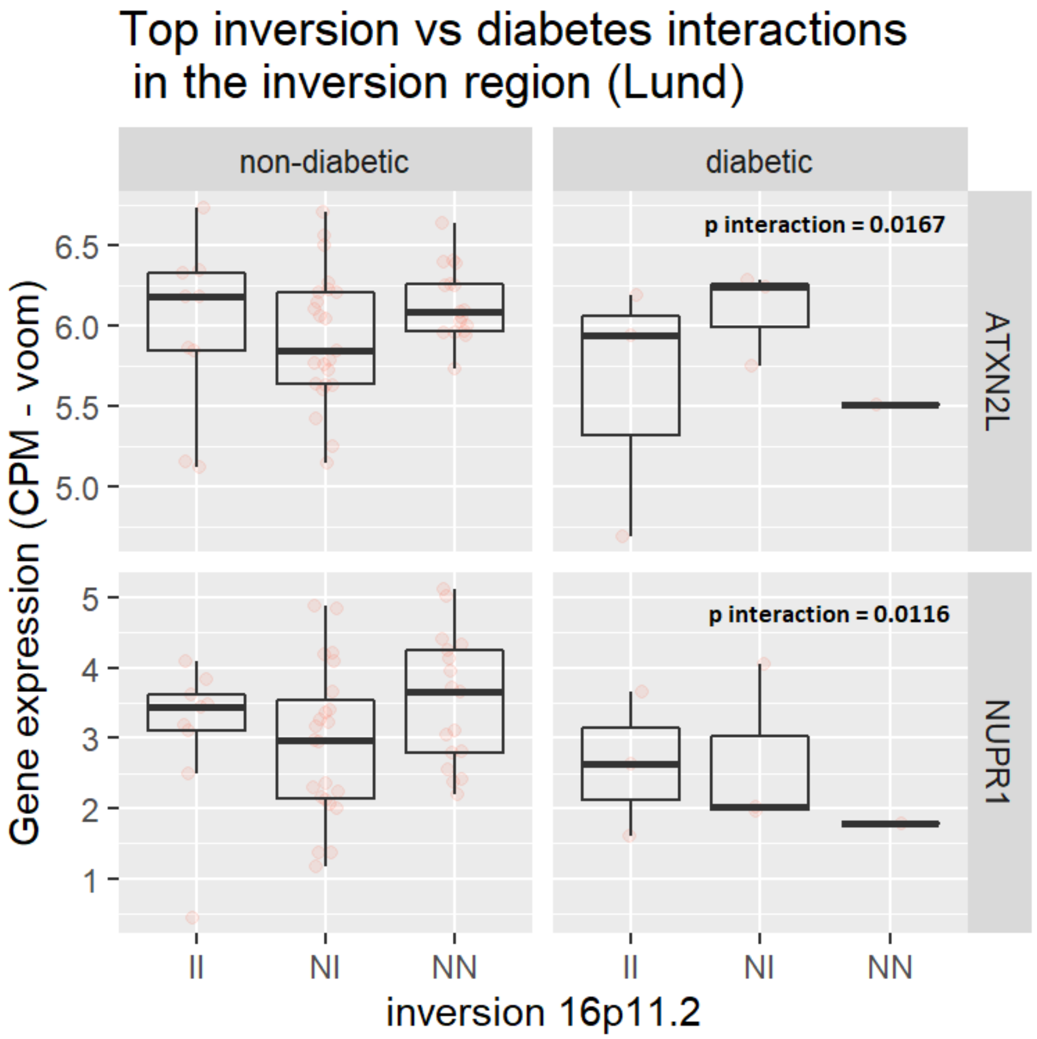


**Figure S6**. **Transcriptomic effect of the interaction between inversion 16p11.2 and diabetes.** Gene expression of the top-leading genes after assessing the interaction between diabetes and inversion at 16p11.2 in 118 pancreatic human islet samples using RNA-sequencing and high-density genotyping data (see **Methods**). The analysis shows that diabetic individuals carrying N-allele have lower gene expression.
